# In Pursuit of a Better Broiler: Growth, Efficiency and Mortality of 16 Strains of Broiler Chickens

**DOI:** 10.1101/2020.10.15.341586

**Authors:** Stephanie Torrey, Mohsen Mohammadigheisar, Midian Nascimento dos Santos, Daniel Rothschild, Lauren Dawson, Zhenzhen Liu, Elijah Kiarie, A. Michelle Edwards, Ira Mandell, Niel Karrow, Dan Tulpan, Tina Widowski

## Abstract

To meet the growing consumer demand for chicken meat, the poultry industry has selected broiler chickens for increasing efficiency and breast yield. While this high productivity means affordable and consistent product, it has come at a cost to broiler welfare. There has been increasing advocacy and consumer pressure on primary breeders, producers, processors and retailers to improve the welfare of the billions of chickens processed annually. Several small-scale studies have reported better welfare outcomes for slower growing strains compared to fast growing, conventional strains. However, these studies often housed birds with range access or used strains with vastly different growth rates. Additionally, there may be traits other than growth, such as body conformation, that influence welfare. As the global poultry industries consider the implications of using slower growing strains, there was a need for a comprehensive, multidisciplinary examination of broiler chickens with a wide range of genotypes differing in growth rate and other phenotypic traits. To meet this need, our team designed a study to benchmark data on conventional and slower growing strains of broiler chickens reared in standardized laboratory conditions. Over a two-year period, we studied 7,528 broilers from 16 different genetic strains. In this paper, we compare the growth, efficiency and mortality of broilers to one of two target weights (TW): 2.1 kg (TW1) and 3.2 kg (TW2). We categorized strains by their growth rate to TW2 as conventional (CONV), fastest slow strains (FAST), moderate slow strains (MOD) and slowest slow strains (SLOW). When incubated, hatched, housed, managed and fed the same, the categories of strains differed in body weights, growth rates, feed intake and feed efficiency. At 48 days of age, strains in the CONV category were 835-1264 g heavier than strains in the other categories. By TW2, differences in body weights and feed intake resulted in a 22 to 43-point difference in feed conversion ratios. Categories of strains did not differ in their overall mortality rates.

## INTRODUCTION

As a global protein source, chicken production is more sustainable than beef or pork in terms of carbon emissions (Clune et al., 2017), and with the burgeoning human population and widespread acceptance of chicken meat consumption, chickens began dominating global meat production in 2020. Genetic selection has resulted in a larger, leaner, more efficient chicken that reaches a market weight of 2.1 kg weeks earlier than even 35 years ago (Havenstein et al., 2003; Zuidhof et al., 2014). However, this level of production is associated with animal welfare concerns, as modern commercial strains of broiler chickens are reported to be unable to perform natural and motivated behaviours (Bokkers and Koene, 2004; Bokkers et al., 2007) or may experience lameness (Bassler et al., 2013; Kittelsen et al., 2017). Through genetic selection, the prevalence of some health conditions such as tibial dyschondroplasia (TD), long bone deformities and ascites has been decreasing over the past 30 years (Kapell et al., 2012a, 2012b; Rekaya et al., 2013). Despite these improvements, recent studies have found 15 to 25% of broilers were moderately to severely lame, with lameness increasing with increasing body weight (Opengart et al., 2018; Wilhelmsson et al., 2019). As moderate to severe lameness is considered painful (Nääs et al., 2009; McGeown et al., 2009), this represents a significant welfare concern, particularly as production of heavy broilers (eviscerated weight > 2 kg) has doubled in the past two decades (AAFC, 2019) and average broiler body weights increase every year (National Chicken Council, 2020).

There is growing concern that conventional broiler production will continue to pose welfare concerns for the chickens (e.g., Dawkins and Layton, 2012; Hartcher et al., 2019). As a result, there is increasing attention being paid to the use of alternative genetic strains of “slow” growing broiler chickens. However, there are different definitions of slower growing among global regulations and animal welfare standards. For example, German regulation stipulates that slower growing strains must grow at a maximum of 80% of the rate of conventional broilers, which would give an approximate growth rate of 48-50 g/d (Federal Office for Agriculture and Food, Working Group of the Federal States on Organic Farming, 2009). The Label Rouge program in France requires broilers to be in production for at least 81 days to grow to 2.1 kg, leading to a growth rate of 26 g/d (Volaille Label Rouge, 2020). The Dutch Beter Leven (Better Life) program specifies that slower growing broilers are processed at 56 days or later, with a growth rate less than 45 g/d (Beter Leven, 2016). In essence, there is no standard definition of a slower growing broiler, but the alternative strains of broilers may require anywhere between 7 and 50 extra days in production to reach a similar body weight as conventional broiler chickens.

As part of a large, comprehensive project examining the health, welfare and productivity of broiler chickens, we studied the growth and efficiency of different strains of broiler chickens, with the objective to benchmark data on conventional and slower growing strains of broiler chickens reared under standardized laboratory conditions. We studied 16 different genetic strains of broiler chickens over a two-year period to understand differences in behaviour, activity, physiology, anatomy, mortality, growth, feed efficiency and carcass and meat quality as they relate to the strains’ growth rates and age. In this manuscript, we describe the overall project methodology and compare the production, efficiency and mortality of 16 strains of broiler chickens.

## MATERIALS AND METHODS

Over eight trials, 7,528 broiler chickens from 16 strains were hatched and reared at the Arkell Poultry Research Station of the University of Guelph. All procedures used in this experiment were approved by the University of Guelph’s Animal Care Committee (AUP # 3746) and were in accordance with the guidelines outlined by the Canadian Council for Animal Care (NFACC, 2016).

### Incubation, hatch and neonatal processing

Hatching eggs were received from primary breeding companies 1-5 days prior to set date. Breeding flocks were located within North America. Identifying information (e.g., company and genetic strain name) were removed and each case for each strain was relabelled with an alphabetical code (e.g., A, B, C etc.). Experimenters were blind to genetic strain identities for all measurements. Eggs were stored at 15°C until removal for incubation. Individual eggs were labelled with their strain code using pencil and segregated by strain among incubation trays. Eggs were incubated at 37.5°C, 55% RH, with turns every hour, for 18 days. On day 19 of incubation, eggs were candled, and fertile eggs were placed in the hatcher for 3 days at 36.9°C, 65% RH. All unhatched eggs were broken, and live embryos and surplus chicks were euthanized using carbon dioxide. Any unthrifty or deformed chicks were also euthanized.

Individual birds were vent sexed by an experienced technician and weighed. Birds received vaccines to protect against infectious bronchitis virus (Merial bronchitis vaccine, Mass type, live virus spray vaccine; Boehringer Ingelheim Animal Health Canada Inc. Burlington, ON, Canada), Marek’s Virus Disease (Vectormune^®^ IBD+ Rispens subcutaneous vaccine, Ceva Animal Health, Cambridge, ON, Canada), and coccidiosis (Immucox^®^ 5 spray vaccine, Ceva Animal Health, Cambridge, ON, Canada; spray vaccine).

### Housing

All chickens were reared in a single room, over time, at the Arkell Poultry Research Station, Guelph, ON, Canada. This room contained 28 floor pens, each measuring 160 × 238 cm. Each pen had solid plastic white walls. Pens were equipped with a round pan feeder (diameter = 33.75cm), 5 nipple drinkers, a 25° ramp to a raised platform that was 30 cm above litter, a hanging round scale (diameter = 50.8cm), a mineral PECKstone (cut into ¼ the original size; Protekta, Inc., Lucknow, ON, Canada), and nylon ropes with strips of polyester cloth tied to the end as enrichment (Figure 1). Softwood shavings were used as litter, and litter was replaced at the start of each trial. There were no windows, although two air vents (positioned above pens 9, 10, 19, 20 and 12, 13, 14, 15) allowed in the occasional natural light. On the first three days, 23 hr of light was provided to ensure chicks found food and water (Table 1). Thereafter, 16 hours of light was provided, with one 8-hr dark period. Light intensity was kept at 20 lux via dimmable LED lights (12W, 3000K warm white, flicker-free bulbs with high grade single chip; Think LED, Cambridge, ON, Canada). Temperature was 32°C at placement and was reduced as birds aged (Table 1).

**Figure 1.**
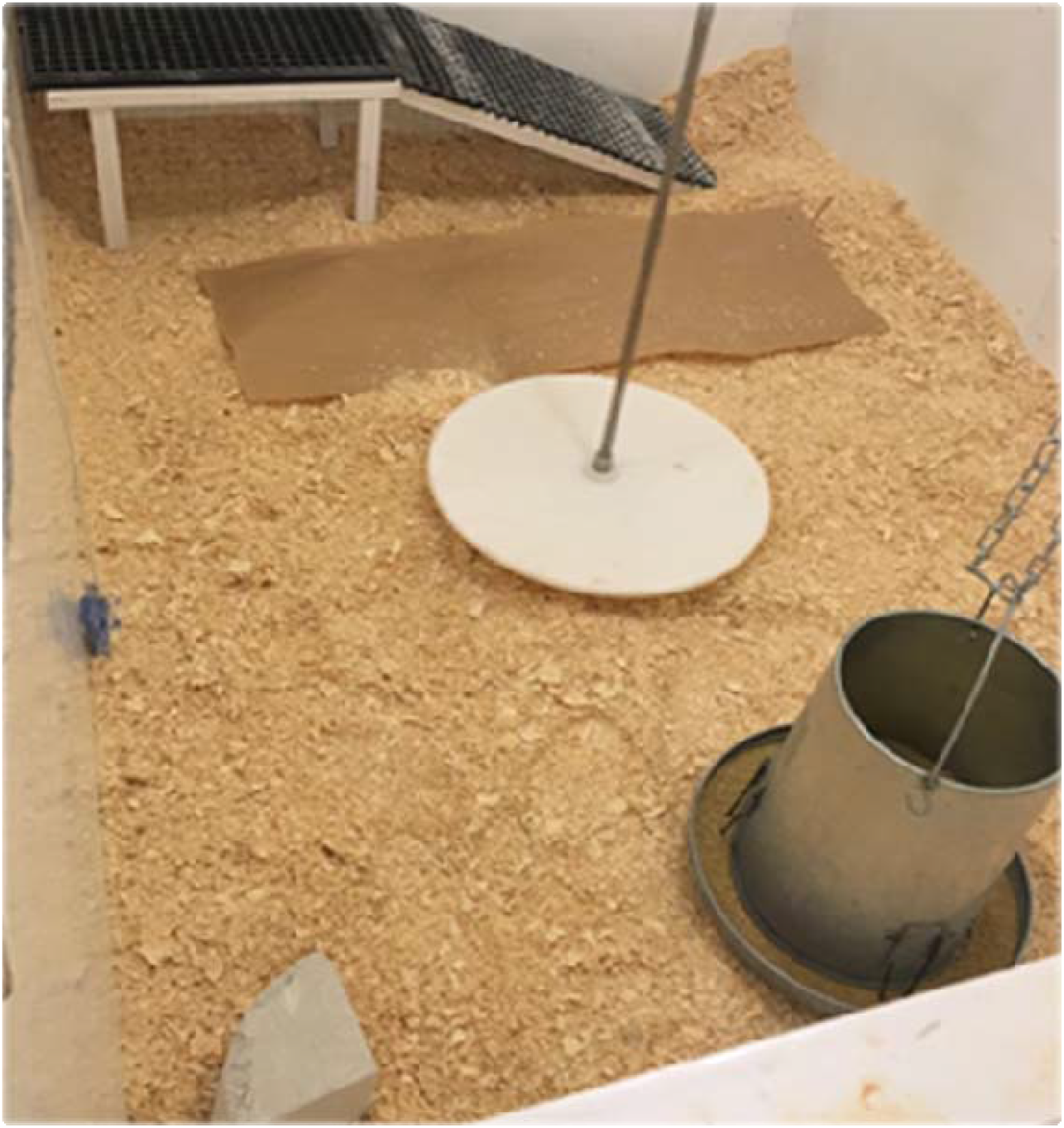
Enrichments within pens. Enrichments provided included a ramp with 25° incline to a platform 30cm above the floor, a hanging scale, PECKStone mineral block (cut into 1/4 of its original size), and hanging (blue) polyester cloth along one wall. Water lines (not shown; they were lowered prior to bird placement) were equipped with five nipple drinkers. Round pan feeders were provided with all vegetarian, antibiotic free feed. Wood shavings were used as litter substrate. Brown paper with scattered feed was provided for the first week of life to aid chicks in finding feed and water. Above the pen walls, netting was affixed to a wooden frame to prevent birds from jumping out of their pens.

**Table 1.** Temperature and lighting schedule through the experiment. A 15-minute dawn/dusk schedule was used, with full lights on at 06:00. Light intensity was maintained at 20 lux.

| Day/Week | Temperature °C | Lighting |
| --- | --- | --- |
| Day 1 | 32 | 23L:1D |
| Day 4 | 32 | 16L:8D |
| Day 5 | 31 | 16L:8D |
| Week 2 | 29 | 16L:8D |
| Week 3 | 27 | 16L:8D |
| Week 4 | 24 | 16L:8D |
| Week 5 | 21 | 16L:8D |

### Diet

An all-vegetable, antibiotic-free diet was provided *ad libitum* (Table 2a). Diets were formulated and milled at the University of Guelph to meet the specifications of a slower growing broiler (Mohammadigheisar et al 2020). Diets were provided in three phases (starter, grower, and finisher). Diets were changed when strains reached the same approximate feed intake as the conventional strain, except for Trial 1 when diets were changed for all birds at 14 and 28 d (onto grower and finisher diets, respectively). In Trial 2, diets were changed based on feed intake, although birds were weighed on days 14 and 28. Therefore, data for growth and efficiency for consumption of equal amounts of feed excludes Trials 1 and 2, whereas data for target weights 1 and 2 include all 8 trials. Each pen transitioned to the next phase diet upon consumption of approximately 500 g/bird starter diet and 1470g/bird grower diet. Diets were a fine crumble for starter phase, coarse crumble for grower phase, and short pellet for finisher stage. Representative samples of feed from each phase were submitted to a commercial laboratory (SGS Inc, Guelph, ON, Canada) for proximate and minerals analyses and AME was estimated based on proximate analyses (Table 2b).

**Table 2a.** Feed formulation and calculated composition for three-phase broiler diets fed to all broilers. Diets were all-vegetable and antibiotic-free, and were formulated for slower growth. Nutrient content represents mean content over eight trials. Approximately 500 g/bird and 1470 g/bird of starter and grower feed, respectively, was provided before an individual pen was switched to the next diet.

| **Ingredients** | **Starter** | **Grower** | **Finisher** |
| --- | --- | --- | --- |
| Corn, Grain | 50.40 | 54.11 | 57.85 |
| Soybean Meal -46% | 28.04 | 25.98 | 21.07 |
| Wheat | 7.13 | 7.44 | 8.77 |
| Corn Gluten Meal-60% | 4.49 | 2.70 | 3.00 |
| Soybean Oil | 3.90 | 4.27 | 4.10 |
| MCP | 1.83 | 1.61 | 1.45 |
| Limestone | 1.65 | 1.48 | 1.36 |
| NaCl | 0.36 | 0.37 | 0.37 |
| NaHCO<sub>3</sub> | 0.29 | 0.20 | 0.21 |
| DL-Methionine | 0.27 | 0.26 | 0.22 |
| L-Lysine HCl, 78% | 0.33 | 0.28 | 0.31 |
| L-Threonine-98% | 0.09 | 0.08 | 0.10 |
| Choline Chloride-60% | 0.22 | 0.22 | 0.20 |
| VT Premix | 1.00 | 1.00 | 1.00 |
| **Calculated composition** |  |  |  |
| Dry Matter, % | 85.03 | 84.56 | 84.63 |
| Crude Protein, % | 21.50 | 19.71 | 18.00 |
| Ca, % | 0.96 | 0.86 | 0.78 |
| Avail. P, % | 0.48 | 0.43 | 0.39 |
| ME, Mcal/kg | 3.04 | 3.09 | 3.13 |
| dMet+Cys, % | 0.86 | 0.80 | 0.73 |
| dMet, % | 0.58 | 0.54 | 0.48 |
| diLys, % | 1.15 | 1.05 | 0.95 |
| dTrp, % | 0.22 | 0.21 | 0.19 |
| dThr, % | 0.75 | 0.69 | 0.64 |
| dIle, % | 0.81 | 0.74 | 0.66 |
| dVal, % | 0.87 | 0.80 | 0.72 |
| dArg, % | 1.22 | 1.14 | 1.00 |
| Na, % | 0.22 | 0.20 | 0.20 |
| Cl, % | 0.28 | 0.28 | 0.28 |

**Table 2b.** Analyzed nutrient composition for three-phase broiler diets fed to all broilers. Diets were all-vegetable and antibiotic-free, and were formulated for slower growth. Nutrient content represents mean content over eight trials.

| Analyzed nutrient content | Starter | Grower | Finisher |
| --- | --- | --- | --- |
| Metabolizable Energy, mcal/kg | 2.874 | 2.98 | 2.91 |
| Moisture, % | 12.32 | 11.88 | 11.71 |
| Dry matter, % | 87.68 | 88.12 | 88.29 |
| Crude protein (N x 6.25), % | 21.46 | 20.50 | 19.58 |
| Calcium, % | 1.03 | 1.02 | 0.89 |
| Total phosphorus, % | 0.77 | 0.71 | 0.65 |
| Potassium, % | 0.84 | 0.82 | 0.74 |
| Magnesium, % | 0.16 | 0.16 | 0.15 |
| Sodium, % | 0.21 | 0.20 | 0.21 |
| Crude fat, % | 5.70 | 6.32 | 6.18 |
| Starch, as is | 35.93 | 36.97 | 39.27 |
| Ethanol soluble carbohydrates, as is | 4.48 | 4.17 | 3.72 |

### Experimental design

In total, 16 strains of broiler chickens were used in this study (Table 3): 3 conventional and 13 slower growing strains. Most strains were represented in 4 pens within each of 3 trials, although there were exceptions due to availability or hatchability (Table 4). Pens within the room were blocked into four groups based on location (Figure 2). Within a trial, birds from each strain were placed in one pen in each of the four blocks. Trials ran from April 2018 until November 2019.

**Table 3.** Strains used in benchmarking experiment. Strains are listed by (breeder) estimated days to reach 2.1 kg, estimated average daily gain (ADG, g/d) and realized ADG to the first target weight (approximately 2.1 kg) and the second target weight (approximately 3.2 kg).

| Strain | Estimated days<br>to 2.1 kg | Estimated ADG, g/d | Realized ADG, g/d |  | Category<sup>2</sup> |
| --- | --- | --- | --- | --- | --- |
|  |  |  | Target wt 1 | Target wt 2 |  |
| A<sup>1</sup> | 34 | 61.76 | 49.12 | 62.65 | - |
| B | 36 | 58.33 | 54.03 | 68.70 | CONV |
| C | 37 | 56.76 | 55.26 | 66.01 | CONV |
| M | 50 | 42.00 | 51.97 | 55.46 | FAST |
| F | 43 | 48.84 | 53.08 | 55.29 | FAST |
| I | 45 | 46.67 | 47.10 | 54.65 | FAST |
| G | 44 | 47.73 | 47.40 | 53.54 | FAST |
| H | 44 | 47.73 | 47.86 | 51.22 | MOD |
| E | 42 | 50.00 | 53.27 | 50.83 | MOD |
| S | 51 | 41.18 | 45.57 | 50.61 | MOD |
| O | 40 | 52.50 | 47.78 | 50.15 | MOD |
| J | 47 | 44.68 | 42.73 | 47.73 | SLOW |
| D | 50 | 42.00 | 42.44 | 45.56 | SLOW |
| N | 50 | 42.00 | 39.82 | 44.06 | SLOW |
| K | 49 | 42.86 | 39.31 | 43.58 | SLOW |
| T<sup>1</sup> | 120 | 17.50 | -- | 19.78 | - |

<sup>1</sup> Strains A and T were represented in few pens (4) and were therefore not included in statistical analyses. The data are included for descriptive purposes only.
<sup>2</sup> Strains were divided into categories based on realized Average Daily Gain to Target Weight 2. This facilitated analyses and enables generalizations based on growth rates.

**Table 4.** Placement of strains within trials. Numbers represent number of pens for that strain within the trial. The ideal was 4 pens in each of 3 trials. This did not occur for 5 strains due to availability (A, E, M, T), or hatchability (G, T). Strains E and G ultimately had 12 pens in total due to either more pens in one trial (E) or representation in an additional trial (G).

| Strain | Trial 1 | Trial 2 | Trial 3 | Trial 4 | Trial 5 | Trial 6 | Trial 7 | Trial 8 | Total |
| --- | --- | --- | --- | --- | --- | --- | --- | --- | --- |
| A |  |  | 4 |  |  |  |  |  | 4 |
| B |  | 4 |  | 4 | 4 |  |  |  | 12 |
| C | 4 |  | 4 |  |  |  |  | 4 | 12 |
| D | 4 | 4 | 4 |  |  |  |  |  | 12 |
| E |  | 4 |  |  | 8 |  |  |  | 12 |
| F | 4 |  |  | 4 | 4 |  |  |  | 12 |
| G |  |  |  | 2 | 2 | 4 | 4 |  | 12 |
| H | 4 | 4 | 4 |  |  |  |  |  | 12 |
| I |  |  |  | 4 |  | 4 | 4 |  | 12 |
| J |  |  |  | 4 | 4 | 4 |  |  | 12 |
| K |  |  |  |  |  | 4 | 4 | 4 | 12 |
| M |  | 4 | 4 |  |  |  |  |  | 8 |
| N |  |  |  | 4 | 4 | 4 |  |  | 12 |
| O |  |  |  |  |  | 4 | 4 | 4 | 12 |
| S |  |  |  |  |  | 4 | 4 | 4 | 12 |
| T | 2 |  |  |  |  |  |  | 2 | 4 |

**Figure 2.**
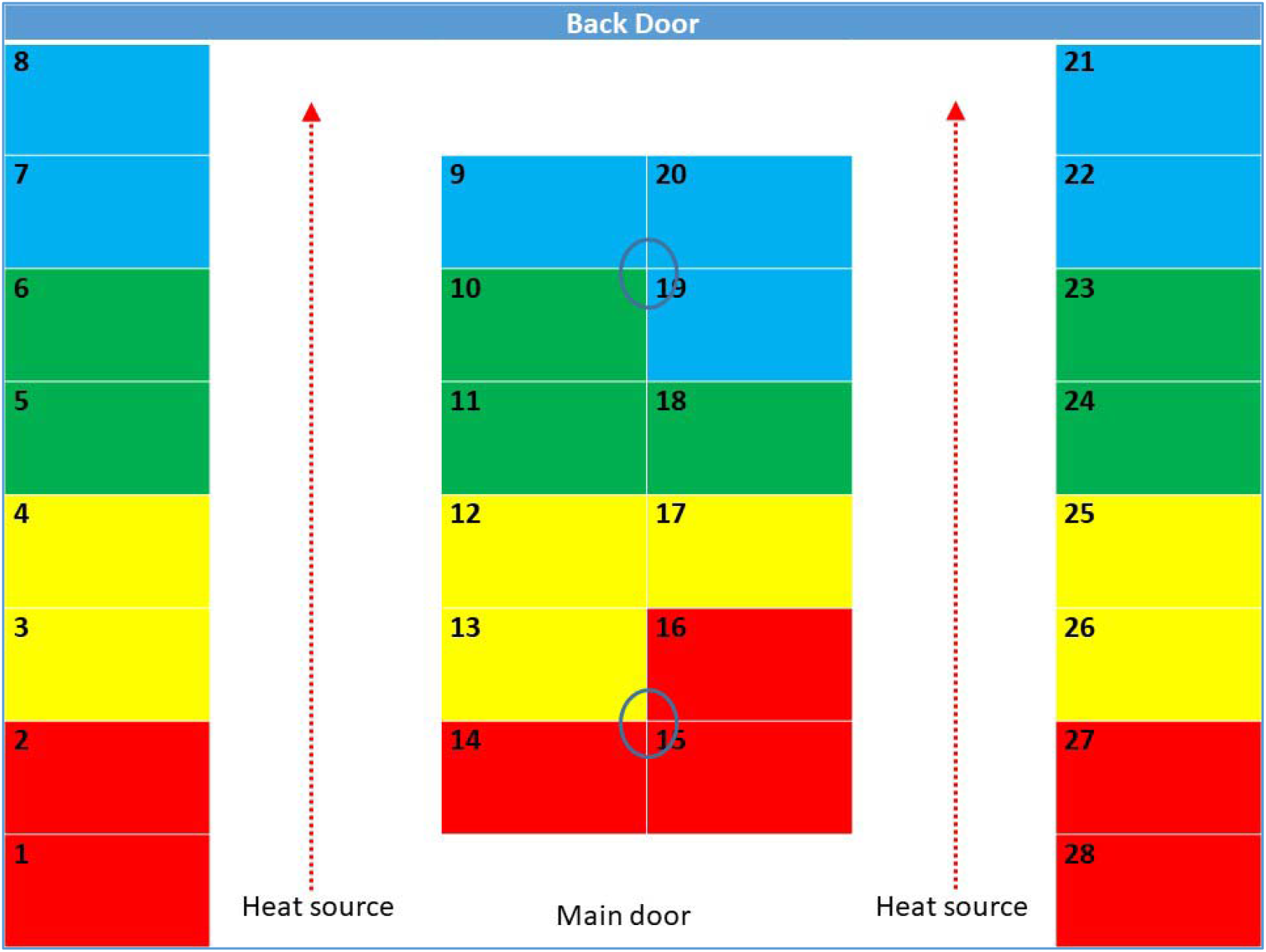
Room layout and block design for trials. Due to the temperature gradient from the main door to the back door and its effect on pen microclimate, four blocks (denoted by different colours) were created within the room, each including seven pens (Block 1: pens 1,2,14,15,16,27,28; Block 2: pens 3,4,12,13,17,25,26; Block 3: pens 5,6,11,18,19,23,24; Block 4: pens 7,8,9,10,20,21,22). Each block included two pens each on the left and right exterior walls, and three pens within the interior of the room. For each trial, each strain used within that trial was represented in each block. The blue circles denote the location of the ventilation outlets over pens 9, 10, 19, 20 and 13, 14, 15, 16, that permitted natural light.

At hatch, all birds of each sex from each strain were weighed as a group, and the average hatch weight for each sex and strain was determined. Then, 22 males and 22 females per strain were selected for each pen based on their average weight being within 10% of their sex and strain’s average body weight. In each pen, 44 birds were placed (22 males and 22 females) on day of hatch.

In Trial 1, half of the birds from each pen (equal numbers of males and females) were processed at 34 d (for strain C) or at 48 d (for other strains), and birds remaining in the two pens per strain were combined to maintain stocking density. In all the other trials, entire pens were processed together to eliminate the effect of mixing birds. On day 34, half of the pens of conventional strains (A, B, C) were processed. The remaining pens of conventional strains were reduced to 38 birds (19 males, 19 females) each. All pens of slower growing strains were reduced to 42 birds (21 males, 21 females) each at 34 d. This was done to maintain as similar a stocking density (kg/m^2^) as possible. On day 48, the remaining pens of conventional strains were processed along with half of the pens with slower growing strains (D-T). The remaining pens of slower growing strains were reduced to 38 birds (19 males, 19 females) and processed on day 62.

To facilitate comparisons, two ‘target weights’ (**TW)** were chosen to represent common processing weights. The first target weight (**TW1)** was estimated to 2.1 kg. Based on the breeder estimated days to 2.1 kg, TW1 was calculated for conventional strains (A, B, C) at 34 d of age and for slower growing strains (D-T) at 48 d of age. The second target weight (**TW2)** was estimated to 3.2 kg. Based on breeder estimated days to 3.2 kg, TW2 was calculated for conventional strains at 48 d of age and for slower growing strains at 62 d of age. These ages were chosen to align with the mean ages for the conventional and slower growing strains, and for convenience in processing at a commercial plant.

### Data collection

Body weights (**BW**) were measured on a group basis on d0 and at each change of diet phase. For conventional strains, this was on d14, d28, d35 and the day prior to processing. The day of diet phase change was not standard among slower growing strains. Group weights were determined on the day prior to processing. Average daily gain (**ADG)** was calculated based on average group weights and was adjusted for mortalities and culls.

Feed intake (**FI)** was determined on a per-pen basis and was determined based on diet phase. Conventional strains (A, B, C) transitioned to different diet phases at d14 and d28. Their feed intake was determined at d14, d28, d35 and on the day of processing. Slower growing strains transitioned to different diet phases depending on their feed intake (500 g/bird of starter diet; 1470 g/bird of grower diet). Feed intake was determined for each diet phase, and until processing day. Feed intake was corrected for mortality as it occurred. Feed conversion ratio (**FCR**) was calculated as grams feed intake / grams body weight, based on average pen feed intake and average pen body weight. **cFCR** was the feed conversion ratio adjusted for mortalities and culls.

Mortality was recorded as it occurred (along with the bird’s body weight and cause of death, if known). Birds were checked at a minimum of once per day and birds were euthanized by manual cervical dislocation if they were lame (e.g., unable to support their body weight or walk more than 1 m), unthrifty (e.g., visibly smaller than pen-mates, ruffled feathers, lethargic), or had serious injuries (e.g., broken wing). Mortality is reported as the percentage of birds culled and found dead.

### Statistical analyses

To aid analyses and enable generalization by growth rate, we categorized strains by their average daily gain to TW2 into one of four categories (conventional, CONV; fastest slow growing, FAST; moderate slow growing, MOD; slowest slow growing, SLOW). CONV encompassed 2 strains (B and C), FAST encompassed 4 strains (F, G, I and M), MOD encompassed 4 strains (E, H, O and S), and SLOW encompassed 4 strains (D, J K and N); two strains (A and T) were excluded from statistical analyses due to low sample sizes (Table 3). All data were input into Excel prior to statistical analyses with SAS^®^ v9.4. Data were analyzed using a generalized linear mixed model (GLIMMIX procedure), with pen as the independent experimental unit. Significance level was set at p-values less than 0.05. Separate models were built to analyze effects according to age and target weight. For both, category and strain nested within category were used as fixed effects, and trial and block were used as random effects. To analyze differences by age, age was fit into a repeated structure that produced the lowest Akaike information criterion value. To analyze differences by target weight, target weight was included as a fixed effect. Trial and block were included in the covariance structure as random effects.

Age was fit into a repeated structure, where appropriate. Contrast statements were used to compare categories, and strains within categories. Pairwise comparisons were adjusted for multiple comparisons using the Tukey adjustment. Model assumptions were assessed using a scatterplot of studentized residuals, linear predictor for linearity, and a Shapiro-Wilk test for normality. The Gaussian distribution was used as the default distribution, but if the assumption for linearity was not met, data were transformed via the selection of a different distribution. Mortality data were analyzed as ordinal (i.e., number of birds per pen that died) since zeros were overrepresented and normality could not be achieved with any transformation. Ordinal data were analyzed using GLIMMIX with a multinomial distribution. Odds ratios were calculated to compare differences in the levels of fixed effects.

## RESULTS

Data are presented first as comparisons between categories, and then as comparisons between strains within categories.

### Production and efficiency through consumption of the same feed allotment by category

Since we categorized strains by growth rate post-hoc (Table 3), BW and ADG differed among categories, with CONV having the highest BW and ADG, and the SLOW having the lowest BW and ADG (Table 5), through the consumption of the same allotment of feed.

**Table 5.** Differences in body weight, growth, feed intake and feed conversion (LS-means ± SEM) between categories of broilers through consumption of similar amounts of starter, grower and finisher rations. The days (mean ± SE) during which each category was fed each phase is shown.

|  | Category |  |  |  |
| --- | --- | --- | --- | --- |
|  | CONV | FAST | MOD | SLOW |
| **Starter** |  |  |  |  |
| BW (kg) | 0.47 ± 0.011<sup>a</sup> | 0.42 ± 0.010<sup>b</sup> | 0.42 ± 0.010<sup>b</sup> | 0.39 ± 0.010<sup>c</sup> |
| ADG (g) | 29.95 ± 0.459<sup>a</sup> | 24.67 ± 0.429<sup>b</sup> | 23.36 ± 0.429<sup>c</sup> | 20.77 ± 0.425<sup>d</sup> |
| ADFI (g) | 36.42 ± 0.577<sup>a</sup> | 31.15 ± 0.532<sup>b</sup> | 30.93 ± 0.533<sup>b</sup> | 28.94 ± 0.527<sup>c</sup> |
| cFCR<sup>2</sup> (g feed/g BW) | 1.22 ± 0.021<sup>c</sup> | 1.26 ± 0.018<sup>c</sup> | 1.33 ± 0.018<sup>b</sup> | 1.40 ± 0.017<sup>a</sup> |
| FCR (g feed/g BW) | 1.22 ± 0.021<sup>c</sup> | 1.26 ± 0.018<sup>c</sup> | 1.33 ± 0.018<sup>b</sup> | 1.40 ± 0.018<sup>a</sup> |
| Days<sup>3</sup> | 14.00 ± 0.00 | 15.47 ± 0.61 | 15.92 ± 0.94 | 16.60 ± 0.87 |
| **Grower** |  |  |  |  |
| BW (kg) | 1.50 ± 0.023<sup>a</sup> | 1.33 ± 0.021<sup>b</sup> | 1.32 ± 0.021<sup>b</sup> | 1.23 ± 0.021<sup>c</sup> |
| ADG (g) | 73.50 ± 1.164<sup>a</sup> | 56.02 ± 1.036<sup>b</sup> | 53.55 ± 1.036<sup>c</sup> | 50.33 ± 1.020<sup>d</sup> |
| ADFI (g) | 105.77 ± 4.400<sup>a</sup> | 86.37 ± 4.342<sup>b</sup> | 82.87 ± 4.344<sup>c</sup> | 83.03 ± 4.336<sup>c</sup> |
| cFCR (g feed/g BW) | 1.41 ± 0.085<sup>c</sup> | 1.54 ± 0.084<sup>b</sup> | 1.55 ± 0.084<sup>b</sup> | 1.65 ± 0.084<sup>a</sup> |
| FCR (g feed/g BW) | 1.41 ± 0.085<sup>c</sup> | 1.54 ± 0.084<sup>b</sup> | 1.56 ± 0.084<sup>b</sup> | 1.65 ± 0.084<sup>a</sup> |
| Days | 14.00 ± 0.00 | 16.56 ± 1.03 | 17.14 ± 0.59 | 17.50 ± 1.88 |
| **Finisher** |  |  |  |  |
| BW (kg) | 2.02 ± 0.026<sup>a</sup> | 1.95 ± 0.020<sup>ab</sup> | 1.94 ± 0.020<sup>b</sup> | 1.77 ± 0.020<sup>c</sup> |
| ADG (g) | 89.12 ± 2.819<sup>a</sup> | 78.61 ± 2.241<sup>b</sup> | 83.94 ± 2.235<sup>ab</sup> | 57.61 ± 2.166<sup>c</sup> |
| ADFI (g) | 159.02 ± 4.345<sup>a</sup> | 145.00 ± 2.382<sup>b</sup> | 158.64 ± 3.375<sup>a</sup> | 123.15 ± 3.261<sup>c</sup> |
| cFCR (g feed/g BW) | 1.82 ± 0.060<sup>b</sup> | 1.86 ± 0.046<sup>b</sup> | 1.89 ± 0.046<sup>b</sup> | 2.15 ± 0.045<sup>a</sup> |
| FCR (g feed/g BW) | 1.88 ± 0.054<sup>b</sup> | 1.88 ± 0.043<sup>b</sup> | 1.89 ± 0.043<sup>b</sup> | 2.13 ± 0.041<sup>a</sup> |
| Days | 6.13 ± 1.09 | 8.08 ± 1.13 | 7.89 ± 1.01 | 9.48 ± 1.38 |
| **Overall** |  |  |  |  |
| ADG, g/bird | 58.48 ± 0.609<sup>a</sup> | 48.27 ± 0.502<sup>b</sup> | 46.81 ± 0.501<sup>c</sup> | 40.23 ± 0.488<sup>d</sup> |
| ADFI, g/bird | 87.19 ± 2.084<sup>a</sup> | 76.88 ± 2.017<sup>b</sup> | 75.62 ± 2.019<sup>b</sup> | 70.53 ± 2.010<sup>c</sup> |
| cFCR, g feed/g BW | 1.49 ± 0.035<sup>c</sup> | 1.59 ± 0.034<sup>b</sup> | 1.62 ± 0.034<sup>b</sup> | 1.75 ± 0.033<sup>a</sup> |
| FCR, g feed/g BW | 1.51 ± 0.037<sup>c</sup> | 1.60 ± 0.035<sup>b</sup> | 1.62 ± 0.035<sup>b</sup> | 1.75 ± 0.035<sup>a</sup> |
| Age, d | 34.13 ± 0.27 | 40.11 ± 0.27 | 40.94 ± 0.26 | 43.58 ± 0.27 |

<sup>a#b#c</sup> Within a row, values without a common superscript differ, adjP<0.05
<sup>1</sup>Because of differences in methodologies, trials 1-2 were omitted from this data set as diets were changed at 14 and 28 days for all strains.
<sup>2</sup> Mortality adjusted feed conversion ratio.
<sup>3</sup>Mean number of days to consume approximately 0.50 kg starter, 1.48 kg grower and 1.00 kg finisher feed allotment.

Differences in BW (F_3,109_=81.44; P<0.0001) and ADG (F_3,109_=350.08; P<0.0001) among categories were apparent through consumption of the 0.5 kg starter allotment (Table 5). CONV finished their starter feed allotment 1.47 to 2.60 days earlier and had heavier BW and ADG than other categories (Table 5). SLOW had the lowest BW and ADG through consumption of the starter allotment. FAST and MOD did not differ in BW through the starter period. Average daily feed intake (ADFI; F_3,109_=96.05; P<0.0001) and feed conversion ratios (corrected for mortality; cFCR; F_3,109_=36.65; P<0.0001) differed by category through consumption of the starter feed allotment. CONV had the highest, and SLOW had the lowest ADFI. CONV and FAST had similar cFCR and were more efficient than MOD and SLOW. MOD also had better cFCR than SLOW.

CONV consumed 1.48 kg of grower ration in 14 days (Table 5). Other categories needed 2.56 to 3.50 extra days to consume a similar amount. Categories differed in BW (F_3,109_=89.76; P<0.0001), ADG (F_3,109_=211.76; P<0.0001), ADFI (F_3,109_=196.24; P<0.0001) and cFCR (F_3,109_=7.20; P=0.0002), with CONV having heavier BW, higher ADG, higher ADFI and better cFCR than all other categories (Table 5).

Through consumption of approximately 1.0 kg finisher feed, categories differed in their BW (F_3,109_=35.78; P<0.0001), ADG (F_3,109_=37.12; P=<0.0001) and ADFI (F_3,109_=48.32; P<0.0001). CONV was heavier than MOD and SLOW (with FAST intermediate between CONV and MOD), and had higher ADG than other categories (Table 5). CONV and MOD had similar ADFI during this phase, with higher daily feed intake than FAST and SLOW. cFCR differed by category (F_3,109_=6.18; P=0.0006). CONV, FAST and MOD had similar FCRs, and were more feed efficient than SLOW.

### Production and efficiency through consumption of the same feed allotment by strains within category

Within categories, strains differed in BW (F_10,109_=11.25; P<0.0001), ADG (F_10,109_=15.99; P<0.0001), ADFI (F_10,109_=10.26; P<0.0001) and cFCR (F_10,109_=2.95; P=0.0026) during the starter period. During the grower period, BW (F_10,109_=5.54; P<0.0001), ADG (F_10,109_=11.12; P<0.0001) and ADFI (F_10,109_=13.85; P<0.0001) differed between strains within categories. There was no effect of strain within category on cFCR during the growing phase (F_10,109_=1.33; P=0.22). During consumption of the same amount of finisher feed, strains within categories differed in BW (F_10,109_=5.35; P<0.0001), ADG (F_10,109_=4.25; P<0.0001), ADFI (F_10,109_=14.98; P<0.0001) and cFCR (F_10,109_=2.43; P=0.0118).

In the CONV category, strain B was heavier in each dietary phase, and had higher ADG and ADFI during the starter and grower periods, than strain C (Table 6). Both conventional strains had similar FCR in each dietary phase. Although not statistically analyzed, strain A performed similar to strain C in terms of BW, ADG and ADFI.

**Table 6.** Differences in the growth performance (LS-means ± SEM) among CONV strains of broiler chickens through consumption of equal amounts of starter, grower and finisher rations. The days during which each strain was fed each phase is shown.

|  | Strain |  |  |
| --- | --- | --- | --- |
|  | A<sup>2</sup> | B | C |
| **Starter** |  |  |  |
| BW (kg) | 0.45 ± 0.007 | 0.50 ± 0.012<sup>a</sup> | 0.45 ± 0.012<sup>b</sup> |
| ADG (g) | 28.82 ± 0.476 | 31.80 ± 0.550<sup>a</sup> | 28.10 ± 0.572<sup>b</sup> |
| ADFI (g) | 34.10 ± 0.461 | 38.84 ± 0.705<sup>a</sup> | 34.00 ± 0.733<sup>b</sup> |
| cFCR<sup>3</sup> (g feed/g BW) | 1.18 ± 0.013 | 1.23 ± 0.030 | 1.21 ± 0.030 |
| FCR (g feed/g BW) | 1.18 ± 0.013 | 1.23 ± 0.029 | 1.21 ± 0.029 |
| Days<sup>4</sup> | 14.00 ± 0.00 | 14.00 ± 0.00 | 14.00 ± 0.00 |
| **Grower** |  |  |  |
| BW (kg) | 1.36 ± 0.006 | 1.58 ± 0.029<sup>a</sup> | 1.42 ± 0.030<sup>b</sup> |
| ADG (g) | 64.70 ± 0.785 | 77.22 ± 1.499<sup>a</sup> | 69.79 ± 1.553<sup>b</sup> |
| ADFI (g) | 103.68 ± 0.771 | 109.12 ± 4.610<sup>a</sup> | 102.41 ± 4.685<sup>b</sup> |
| cFCR (g feed/g BW) | 1.60 ± 0.026 | 1.41 ± 0.089 | 1.41 ± 0.091 |
| FCR (g feed/g BW) | 1.60 ± 0.026 | 1.41 ± 0.090 | 1.41 ± 0.091 |
| Days | 14.00 ± 0.00 | 14.00 ± 0.00 | 14.00 ± 0.00 |
| **Finisher** |  |  |  |
| BW (kg) | 1.87 ± 0.097 | 2.14 ± 0.036<sup>a</sup> | 1.90 ± 0.036<sup>b</sup> |
| ADG (g) | 93.88 ± 5.024 | 90.62 ± 3.940 | 87.61 ± 3.976 |
| ADFI (g) | 156.10 ± 7.725 | 167.07 ± 6.109 | 150.97 ± 6.141 |
| cFCR (g feed/g BW) | 1.67 ± 0.084 | 1.85 ± 0.084 | 1.79 ± 0.084 |
| FCR (g feed/g BW) | 1.72 ± 0.079 | 1.94 ± 0.076 | 1.83 ± 0.076 |
| Days | 5.50 ± 0.87 | 6.50 ± 0.19 | 5.75 ± 0.49 |
| **Overall** |  |  |  |
| ADG (g) | 54.73 ± 1.839 | 61.28 ± 0.838<sup>a</sup> | 55.68 ± 0.853<sup>b</sup> |
| ADFI (g) | 83.41 ± 2.819 | 91.83 ± 2.309<sup>a</sup> | 82.54 ± 2.379<sup>b</sup> |
| cFCR (g feed/g BW) | 1.52 ± 0.006 | 1.50 ± 0.041 | 1.48 ± 0.043 |
| FCR (g feed/g BW) | 1.54 ± 0.011 | 1.52 ± 0.042 | 1.49 ± 0.044 |
| Age, d | 33.50 ± 0.87 | 34.50 ± 0.19 | 33.75 ± 0.49 |

<sup>a</sup>/<sup>b</sup>/<sup>c</sup> Within a row, values without a common superscript differ, adjP<0.05
<sup>1</sup>Due to differences in methodologies, trials 1-2 were omitted from this data set as diets were changed at 14 and 28 days for all strains.
<sup>2</sup>Because A was represented in a single trial, data for this strain are included for descriptive purposes only and were not subject to statistical analyses.
<sup>3</sup>Mortality adjusted feed conversion ratio.
<sup>4</sup>Mean number of days required to consume approximately 0.50 kg starter, 1.48 kg grower and 1.00 kg finisher feed allotment.

Among the FAST strains, strain M had the heaviest BW and the highest ADG during the starter period (Table 7). They also had higher ADFI than strain G during the same period (Table 7), and higher ADFI than all other FAST strains during the grower period. During the finisher diet phase, strain M had a lighter BW than strain G, and a lower ADG than strain F (Table 7). During the finisher period, strain F had the highest ADG. There were differences in ADFI among FAST strains, with strain M having the highest ADFI in the starter and grower phases, while strains F and I had the highest ADFI for the finisher period (Table 7). Strain M had worse FCR through the consumption of similar amounts of feed compared to the other FAST strains.

**Table 7.** Differences in the growth performance (LS-means ± SEM) among FAST strains of broiler chickens through consumption of equal amounts of starter, grower and finisher rations. The days during which each strain was fed each phase is shown.

|  | Strain |  |  |  |
| --- | --- | --- | --- | --- |
|  | F | G | I | M |
| **Starter** |  |  |  |  |
| BW (kg) | 0.42 ± 0.012<sup>b</sup> | 0.40 ± 0.011<sup>b</sup> | 0.41 ± 0.011<sup>b</sup> | 0.46 ± 0.016<sup>a</sup> |
| ADG (g) | 24.75 ± 0.550<sup>ab</sup> | 23.35 ± 0.488<sup>c</sup> | 23.82 ± 0.490<sup>bc</sup> | 26.75 ± 0.771<sup>a</sup> |
| ADFI (g) | 31.22 ± 0.705<sup>ab</sup> | 29.95 ± 0.617<sup>b</sup> | 30.10 ± 0.621<sup>ab</sup> | 33.31 ± 1.004<sup>a</sup> |
| cFCR<sup>2</sup> (g feed/g BW) | 1.26 ± 0.028 | 1.29 ± 0.023 | 1.27 ± 0.024 | 1.24 ± 0.041 |
| FCR (g feed/g BW) | 1.27 ± 0.029 | 1.29 ± 0.024 | 1.27 ± 0.024 | 1.24 ± 0.042 |
| Days<sup>3</sup> | 15.00 ± 0.00 | 15.83 ± 0.21 | 15.58 ± 0.15 | 15.00 ± 0.00 |
| **Grower** |  |  |  |  |
| BW (kg) | 1.37 ± 0.029 | 1.33 ± 0.025 | 1.34 ± 0.025 | 1.29 ± 0.041 |
| ADG (g) | 54.87 ± 1.499 | 55.44 ± 1.270 | 54.19 ± 1.280 | 59.57 ± 2.184 |
| ADFI (g) | 80.06 ± 4.610<sup>b</sup> | 83.64 ± 4.466<sup>b</sup> | 82.50 ± 4.471<sup>b</sup> | 99.28 ± 5.298<sup>a</sup> |
| cFCR (g feed/g BW) | 1.48 ± 0.089 | 1.53 ± 0.086 | 1.54 ± 0.086 | 1.61 ± 0.103 |
| FCR (g feed/g BW) | 1.47 ± 0.090 | 1.53 ± 0.087 | 1.54 ± 0.087 | 1.61 ± 0.104 |
| Days | 17.00 ± 0.00 | 16.83 ± 0.21 | 16.83 ± 0.11 | 14.00 ± 0.00 |
| **Finisher** |  |  |  |  |
| BW (kg) | 1.97 ± 0.036<sup>ab</sup> | 2.02 ± 0.029<sup>a</sup> | 1.98 ± 0.029<sup>ab</sup> | 1.83 ± 0.051<sup>b</sup> |
| ADG (g) | 87.97 ± 3.940<sup>a</sup> | 78.58 ± 3.193<sup>ab</sup> | 82.51 ± 3.224<sup>ab</sup> | 65.37 ± 5.665<sup>b</sup> |
| ADFI (g) | 157.22 ± 6.109<sup>a</sup> | 135.74 ± 4.946<sup>b</sup> | 151.46 ± 4.989<sup>a</sup> | 135.59 ± 8.729<sup>ab</sup> |
| cFCR (g feed/g BW) | 1.79 ± 0.084 | 1.75 ± 0.068 | 1.85 ± 0.068 | 2.06 ± 0.120 |
| FCR (g feed/g BW) | 1.81 ± 0.076<sup>ab</sup> | 1.75 ± 0.061<sup>b</sup> | 1.85 ± 0.062<sup>ab</sup> | 2.11 ± 0.109<sup>a</sup> |
| Days | 7.00 ± 0.00 | 8.75 ± 0.37 | 7.83 ± 0.24 | 9.00 ± 0.00 |
| **Overall** |  |  |  |  |
| ADG (g) | 49.10 ± 0.838 | 48.24 ± 0.684 | 48.27 ± 0.691 | 47.46 ± 1.218 |
| ADFI (g) | 75.50 ± 2.309<sup>ab</sup> | 73.75 ± 2.155<sup>b</sup> | 75.75 ± 2.160<sup>ab</sup> | 82.54 ± 2.957<sup>a</sup> |
| cFCR (g feed/g BW) | 1.54 ± 0.041<sup>b</sup> | 1.53 ± 0.037<sup>b</sup> | 1.57 ± 0.037<sup>b</sup> | 1.73 ± 0.056<sup>a</sup> |
| FCR (g feed/g BW) | 1.54 ± 0.042<sup>b</sup> | 1.54 ± 0.038<sup>b</sup> | 1.57 ± 0.038<sup>b</sup> | 1.73 ± 0.057<sup>a</sup> |
| Age, d | 39.00 ± 0.00 | 41.42 ± 0.47 | 40.25 ± 0.33 | 38.00 ± 0.00 |

<sup>a#b#c</sup> Within a row, values without a common superscript differ, adjP<0.05
<sup>1</sup> Due to differences in methodologies, trials 1-2 were omitted from this data set as diets were changed at 14 and 28 days for all strains.
<sup>2</sup> Mortality adjusted feed conversion ratio.
<sup>3</sup> Mean number of days to consume approximately 0.50 kg starter, 1.48 kg grower and 1.00 kg finisher feed allotment.

Among the MOD strains, strain E was the heaviest and had the highest ADG through consumption of the starter and grower diets (Table 8). During the starter period, strain S had the lightest BW and lowest ADG and ADFI. During the grower period, strain H had a lower ADG than the other three strains. There was no difference among strains in BW at the end of the finisher allotment. However, there were differences ADG and ADFI; strain H had higher ADG and ADFI than other strains, and strain D had higher ADFI than strains O and S. Through consumption of the three diets, strain E had a higher ADG than strains H and S, and higher ADFI than strains O and S. Strain H had the worst cFCR (Table 8).

**Table 8.** Differences in the growth performance (LS-means ± SEM) among MOD strains of broiler chickens through consumption of equal amounts of starter, grower and finisher rations. The days during which each strain was fed each phase is shown.

|  | Strain |  |  |  |
| --- | --- | --- | --- | --- |
|  | E | H | O | S |
| **Starter** |  |  |  |  |
| BW (kg) | 0.44 ± 0.012<sup>a</sup> | 0.46 ± 0.016<sup>a</sup> | 0.40 ± 0.011<sup>b</sup> | 0.38 ± 0.011<sup>c</sup> |
| ADG (g) | 25.32 ± 0.574<sup>a</sup> | 23.57 ± 0.771<sup>ab</sup> | 23.16 ± 0.489<sup>b</sup> | 21.40 ± 0.489<sup>c</sup> |
| ADFI (g) | 32.72 ± 0.739<sup>a</sup> | 31.92 ± 1.004<sup>ab</sup> | 30.33 ± 0.620<sup>b</sup> | 28.74 ± 0.620<sup>c</sup> |
| cFCR<sup>2</sup> (g feed/g BW) | 1.30 ± 0.030 | 1.35 ± 0.041 | 1.31 ± 0.024 | 1.34 ± 0.024 |
| FCR (g feed/g BW) | 1.30 ± 0.030 | 1.35 ± 0.042 | 1.31 ± 0.024 | 1.34 ± 0.024 |
| Days<sup>3</sup> | 15.00 ± 0.00 | 17.00 ± 0.00 | 15.83 ± 0.24 | 16.25 ± 0.28 |
| **Grower** |  |  |  |  |
| BW (kg) | 1.42 ± 0.030<sup>a</sup> | 1.27 ± 0.041<sup>b</sup> | 1.32 ± 0.025<sup>b</sup> | 1.28 ± 0.025<sup>b</sup> |
| ADG (g) | 60.31 ± 1.585<sup>a</sup> | 45.60 ± 2.184<sup>c</sup> | 55.60 ± 1.281<sup>b</sup> | 52.68 ± 1.281<sup>b</sup> |
| ADFI (g) | 87.76 ± 4.669<sup>a</sup> | 75.66 ± 5.298<sup>b</sup> | 85.12 ± 4.463<sup>ab</sup> | 82.92 ± 4.463<sup>ab</sup> |
| cFCR (g feed/g BW) | 1.46 ± 0.090 | 1.62 ± 0.103 | 1.54 ± 0.086 | 1.59 ± 0.086 |
| FCR (g feed/g BW) | 1.47 ± 0.091 | 1.62 ± 0.104 | 1.54 ± 0.087 | 1.59 ± 0.087 |
| Days | 17.00 ± 0.00 | 18.00 ± 0.00 | 16.75 ± 0.18 | 17.33 ± 0.14 |
| **Finisher** |  |  |  |  |
| BW (kg) | 2.00 ± 0.038 | 1.86 ± 0.051 | 1.97 ± 0.029 | 1.91 ± 0.029 |
| ADG (g) | 82.83 ± 4.169<sup>b</sup> | 103.13 ± 5.665<sup>a</sup> | 76.93 ± 3.240<sup>b</sup> | 72.86 ± 3.240<sup>b</sup> |
| ADFI (g) | 158.81 ± 6.423<sup>b</sup> | 206.29 ± 8.729<sup>a</sup> | 137.39 ± 5.008<sup>c</sup> | 132.06 ± 5.008<sup>c</sup> |
| cFCR (g feed/g BW) | 1.92 ± 0.088 | 2.00 ± 0.120 | 1.81 ± 0.069 | 1.82 ± 0.069 |
| FCR (g feed/g BW) | 1.94 ± 0.080 | 2.01 ± 0.109 | 1.80 ± 0.062 | 1.82 ± 0.062 |
| Days | 7.00 ± 0.00 | 6.00 ± 0.00 | 8.42 ± 0.15 | 8.58 ± 0.15 |
| **Overall** |  |  |  |  |
| ADG (g) | 50.40 ± 0.694<sup>a</sup> | 44.18 ± 1.218<sup>b</sup> | 47.67 ± 0.694<sup>ab</sup> | 45.00 ± 0.694<sup>b</sup> |
| ADFI (g) | 79.15 ± 2.370<sup>a</sup> | 77.51 ± 2.957<sup>ab</sup> | 74.20 ± 2.154<sup>b</sup> | 71.60 ± 2.154<sup>b</sup> |
| cFCR (g feed/g BW) | 1.57 ± 0.043<sup>b</sup> | 1.74 ± 0.056<sup>a</sup> | 1.56 ± 0.037<sup>b</sup> | 1.59 ± 0.037<sup>b</sup> |
| FCR (g feed/g BW) | 1.58 ± 0.044<sup>ab</sup> | 1.74 ± 0.057<sup>a</sup> | 1.56 ± 0.038<sup>b</sup> | 1.59 ± 0.038<sup>ab</sup> |
| Age, d | 39.00 ± 0.00 | 44.00 ± 0.00 | 41.00 ± 0.33 | 42.17 ± 0.41 |

<sup>a,b,c</sup> Within a row, values without a common superscript differ, adjP<0.05
<sup>1</sup> Due to differences in methodologies, trials 1-2 were omitted from this data set as diets were changed at 14 and 28 days for all strains.
<sup>2</sup> Mortality adjusted feed conversion ratio.
<sup>3</sup> Mean number of days to consume approximately 0.50 kg Starter, 1.48 kg Grower and 1.00 kg Finisher feed allotment.

In the SLOW category, there were differences for growth and efficiency variables through consumption of the same feed allotment (Table 9). During the starter and grower periods, strains D and J generally had higher BW, ADG and ADFI than strains K and N. by the finisher period, strain J was heavier than strains D and N. Strain D had the worst feed efficiency during both the starter and finisher periods.

**Table 9.** Differences in the growth performance (LS-means ± SEM) among SLOW strains of broiler chickens through consumption of equal amounts of Starter, Grower and Finisher rations. The days during which each strain was fed each phase is shown.

|  | Strain |  |  |  |
| --- | --- | --- | --- | --- |
|  | D | J | K | N |
| **Starter** |  |  |  |  |
| BW (kg) | 0.41 ± 0.016<sup>a</sup> | 0.40 ± 0.011<sup>a</sup> | 0.37 ± 0.011<sup>b</sup> | 0.37 ± 0.011<sup>b</sup> |
| ADG (g) | 20.58 ± 0.771<sup>ab</sup> | 22.20 ± 0.498<sup>a</sup> | 20.08 ± 0.189<sup>b</sup> | 20.22 ± 0.498<sup>b</sup> |
| ADFI (g) | 31.92 ± 1.004<sup>a</sup> | 28.83 ± 0.632<sup>b</sup> | 27.32 ± 0.620<sup>b</sup> | 27.69 ± 0.632<sup>b</sup> |
| cFCR<sup>2</sup> (g feed/g BW) | 1.55 ± 0.041<sup>a</sup> | 1.30 ± 0.024<sup>c</sup> | 1.36 ± 0.024<sup>bc</sup> | 1.37 ± 0.024<sup>b</sup> |
| FCR (g feed/g BW) | 1.55 ± 0.042<sup>a</sup> | 1.30 ± 0.024<sup>c</sup> | 1.36 ± 0.024<sup>bc</sup> | 1.39 ± 0.024<sup>b</sup> |
| Days<sup>3</sup> | 17.00 ± 0.00 | 16.33 ± 0.14 | 17.00 ± 0.39 | 16.33 ± 0.14 |
| **Grower** |  |  |  |  |
| BW (kg) | 1.22 ± 0.041<sup>ab</sup> | 1.27 ± 0.026<sup>a</sup> | 1.25 ± 0.025<sup>ab</sup> | 1.19 ± 0.026<sup>b</sup> |
| ADG (g) | 54.73 ± 2.184<sup>a</sup> | 51.64 ± 1.304<sup>a</sup> | 46.86 ± 1.281<sup>b</sup> | 48.11 ± 1.304<sup>b</sup> |
| ADFI (g) | 89.20 ± 5.298<sup>a</sup> | 85.24 ± 4.493<sup>a</sup> | 76.66 ± 4.463<sup>b</sup> | 81.02 ± 4.493<sup>ab</sup> |
| cFCR (g feed/g BW) | 1.58 ± 0.103 | 1.67 ± 0.087 | 1.63 ± 0.086 | 1.71 ± 0.087 |
| FCR (g feed/g BW) | 1.58 ± 0.104 | 1.67 ± 0.087 | 1.63 ± 0.087 | 1.71 ± 0.087 |
| Days | 15.00 ± 0.00 | 16.92 ± 0.42 | 19.25 ± 0.35 | 17.17 ± 0.47 |
| **Finisher** |  |  |  |  |
| BW (kg) | 1.69 ± 0.051<sup>b</sup> | 1.85 ± 0.029<sup>a</sup> | 1.83 ± 0.029<sup>ab</sup> | 1.71 ± 0.029<sup>b</sup> |
| ADG (g) | 55.18 ± 5.665 | 59.04 ± 3.251 | 62.38 ± 3.240 | 53.85 ± 3.251 |
| ADFI (g) | 140.15 ± 8.73<sup>a</sup> | 117.16 ± 5.02<sup>ab</sup> | 121.77 ± 5.01<sup>a</sup> | 113.15 ± 5.02<sup>b</sup> |
| cFCR (g feed/g BW) | 2.52 ± 0.120<sup>a</sup> | 1.97 ± 0.069<sup>b</sup> | 2.02 ± 0.069<sup>b</sup> | 2.09 ± 0.069<sup>b</sup> |
| FCR (g feed/g BW) | 2.53 ± 0.109<sup>a</sup> | 1.95 ± 0.062<sup>b</sup> | 1.95 ± 0.062<sup>b</sup> | 2.10 ± 0.062<sup>b</sup> |
| Days | 9.00 ± 0.00 | 9.75 ± 0.51 | 9.17 ± 0.11 | 9.67 ± 0.51 |
| **Overall** |  |  |  |  |
| ADG (g) | 40.13 ± 1.218<sup>ab</sup> | 42.00 ± 0.700<sup>a</sup> | 40.14 ± 0.694<sup>ab</sup> | 38.66 ± 0.700<sup>b</sup> |
| ADFI (g) | 77.37 ± 2.957<sup>a</sup> | 70.46 ± 2.183<sup>ab</sup> | 66.75 ± 2.154<sup>c</sup> | 67.55 ± 2.183<sup>bc</sup> |
| cFCR (g feed/g BW) | 1.91 ± 0.057<sup>a</sup> | 1.68 ± 0.039<sup>b</sup> | 1.65 ± 0.038<sup>b</sup> | 1.76 ± 0.039<sup>a</sup> |
| FCR (g feed/g BW) | 1.91 ± 0.056<sup>a</sup> | 1.69 ± 0.038<sup>c</sup> | 1.66 ± 0.037<sup>c</sup> | 1.76 ± 0.038<sup>b</sup> |
| Age, d | 41.00 ± 0.00 | 43.00 ± 0.28 | 45.42 ± 0.38 | 43.17 ± 0.32 |

<sup>a/b/c</sup> Within a row, values without a common superscript differ, adjP<0.05
<sup>1</sup> Due to differences in methodologies, trials 1-2 were omitted from this data set as diets were changed at 14 and 28 days for all strains.
<sup>2</sup> Mortality adjusted feed conversion ratio.
<sup>3</sup> Mean number of days to consume approximately 0.50 kg Starter, 1.48 kg Grower and 1.00 kg Finisher feed allotment

### Production and efficiency to TW1 and TW2 by category

Through processing, BW differed by category (F_3,106_=53.63; P<0.0001) and target weight (F_1,106_=382.47; P<0.0001), and there was a category by target weight interaction (F_3,106_=14.13; P<0.0001). At TW1, FAST and MOD had higher BW than SLOW and CONV (Table 10). At TW2, FAST were heavier than MOD and SLOW, and both CONV and MOD were heavier than SLOW (Table 10). There was no effect of strain within category (F_10,106_=1.58; P=0.12) nor a target weight by strain within category interaction (F_10,106_=1.08; P=0.38).

**Table 10.** Differences between and within categories in live performance traits (back transformed LS-means ± SEM) at Target Weight 1 (TW1) and Target Weight 2 (TW2). At TW1, CONV was 34 d of age and the other categories were 48 d of age. At TW2, CONV was 48 d of age and the other categories were 62 d of age.

| Cat. | Strain | BW (kg) |  | ADG (g) |  | ADFI (g) |  | cFCR (g feed/g BW) |  |
| --- | --- | --- | --- | --- | --- | --- | --- | --- | --- |
|  |  | TW1 | TW2 | TW1 | TW2 | TW1 | TW2 | TW1 | TW2 |
| CONV | | 1.838 ± 0.0674<sup>b</sup> | 3.202 ± 0.0674<sup>ab</sup> | 55.86 ± 1.245<sup>a</sup> | 68.92 ± 1.246<sup>a</sup> | 87.19 ± 2.084<sup>a</sup> | 102.90 ± 2.498<sup>a</sup> | 1.49 ± 0.035<sup>a</sup> | 1.54 ± 0.061<sup>a</sup> |
| FAST | | 2.367 ± 0.0567<sup>a</sup> | 3.433 ± 0.0567<sup>a</sup> | 51.38 ± 1.004<sup>b</sup> | 56.03 ± 1.004<sup>b</sup> | 83.78 ± 2.201<sup>b</sup> | 97.54 ± 2.201<sup>ab</sup> | 1.70 ± 0.048<sup>bc</sup> | 1.76 ± 0.048<sup>b</sup> |
| MOD | | 2.340 ± 0.0547<sup>a</sup> | 3.170 ± 0.0559<sup>b</sup> | 50.27 ± 0.948<sup>b</sup> | 51.79 ± 0.986<sup>c</sup> | 80.39 ± 2.179<sup>b</sup> | 96.36 ± 2.180<sup>b</sup> | 1.62 ± 0.047<sup>b</sup> | 1.89 ± 0.047<sup>c</sup> |
| SLOW | | 1.938 ± 0.0554<sup>b</sup> | 2.813 ± 0.0549<sup>c</sup> | 41.85 ± 0.965<sup>c</sup> | 46.13 ± 0.951<sup>d</sup> | 74.34 ± 2.389<sup>c</sup> | 89.93 ± 2.168<sup>c</sup> | 1.82 ± 0.056<sup>c</sup> | 1.97 ± 0.046<sup>c</sup> |
| CONV | A<sup>1</sup> | 1.675 ± 0.0181 | 3.068 ± 0.0781 | 50.76 ± 0.549 | 65.28 ± 1.683 | 83.41 ± 2.819 | 101.37 ± 2.492 | 1.52 ± 0.006 | 1.58 ± 0.002 |
| | B | 1.789 ± 0.0819 | 3.316 ± 0.0819 | 55.15 ± 1.737 | 70.48 ± 1.737 | 91.83 ± 2.309 | 107.80 ± 2.992 | 1.50 ± 0.041 | 1.57 ± 0.079 |
| | C | 1.872 ± 0.0818 | 3.110 ± 0.0818 | 56.58 ± 1.736 | 67.35 ± 1.736 | 82.54 ± 2.379 | 98.02 ± 3.169 | 1.48 ± 0.043 | 1.50 ± 0.086 |
| FAST | F | 2.483 ± 0.0814 | 3.446 ± 0.0813 | 53.75 ± 1.727 | 56.08 ± 1.727 | 84.26 ± 2.992 | 99.16 ± 2.992 | 1.60 ± 0.079 | 1.80 ± 0.079 |
| | G | 2.290 ± 0.0821 | 3.343 ± 0.0831 | 48.77 ± 1.741 | 54.76 ± 1.741 | 81.72 ± 3.004 | 97.14 ± 3.004 | 1.74 ± 0.079 | 1.77 ± 0.079 |
| | I | 2.269 ± 0.0822 | 3.424 ± 0.0822 | 48.27 ± 1.744 | 56.08 ± 1.744 | 84.44 ± 3.031 | 95.05 ± 3.031 | 1.82 ± 0.080 | 1.69 ± 0.080 |
| | M | 2.533 ± 0.1000 | 3.489 ± 0.1002 | 54.73 ± 2.123 | 57.18 ± 2.127 | 84.68 ± 3.554 | 98.81 ± 3.554 | 1.64 ± 0.097 | 1.78 ± 0.097 |
| MOD | E | 2.554 ± 0.0849 | 3.157 ± 0.0849 | 54.72 ± 1.800 | 51.69 ± 1.800 | 88.35 ± 3.065<sup>a</sup> | 98.77 ± 3.065<sup>ab</sup> | 1.66 ± 0.082 | 1.97 ± 0.082 |
| | H | 2.248 ± 0.0764 | 3.216 ± 0.0902 | 49.16 ± 1.620 | 52.56 ± 1.913 | 78.10 ± 3.108<sup>ab</sup> | 87.32 ± 3.278<sup>b</sup> | 1.65 ± 0.081 | 1.71 ± 0.088 |
| | O | 2.331 ± 0.0829 | 3.110 ± 0.0829 | 49.59 ± 1.756 | 51.01 ± 1.756 | 80.76 ± 3.093<sup>ab</sup> | 102.1 ± 3.093<sup>a</sup> | 1.64 ± 0.081 | 2.02 ± 0.081 |
| | S | 2.239 ± 0.0829 | 3.165 ± 0.0829 | 47.62 ± 1.756 | 51.91 ± 1.756 | 74.36 ± 3.266<sup>b</sup> | 97.28 ± 3.093<sup>ab</sup> | 1.55 ± 0.087 | 1.86 ± 0.081 |
| SLOW | D | 2.013 ± 0.0825 | 2.873 ± 0.0763 | 43.94 ± 1.750 | 46.70 ± 1.618 | 75.68 ± 3.108 | 91.88 ± 3.108 | 1.80 ± 0.081 | 2.01 ± 0.081 |
| | J | 2.000 ± 0.0820 | 2.943 ± 0.0820 | 42.64 ± 1.740 | 48.20 ± 1.740 | 77.73 ± 3.447 | 90.63 ± 3.000 | 1.80 ± 0.096 | 1.91 ± 0.080 |
| | K | 1.933 ± 0.0829 | 2.782 ± 0.0829 | 41.11 ± 1.756 | 45.63 ± 1.756 | 70.36 ± 4.617 | 88.54 ± 3.093 | 1.83 ± 0.135 | 1.92 ± 0.081 |
| | N | 1.862 ± 0.0820 | 2.688 ± 0.0820 | 39.70 ± 1.740 | 44.00 ± 1.740 | 73.58 ± 3.447 | 88.68 ± 3.000 | 1.87 ± 0.096 | 2.04 ± 0.080 |
| | T<sup>1</sup> | - | 1.271 ± 0.1381 | - | 20.84 ± 0.226 | - | - | - | - |

<sup>1</sup>Because of their small sample size, strains A and T were not included in their category means nor were they included in statistical analyses. Values for these strains are raw means ± standard errors. Data were not collected for strain T at Target Weight 1 due to its low body weight. Feed intake data were unavailable.
<sup>a≠b</sup> Within a column and section, values without a common superscript differ, adjP<0.05.

For ADG, there was a significant effect of category (F_3,106_=103.44; P<0.0001) and a category by target weight interaction (F_4,106_=11.80; P<0.0001; Table 10). At TW1, CONV had the highest ADG and SLOW had the lowest ADG (Table 10). FAST and MOD were intermediate and did not differ from each other (Table 10). At TW2, CONV had the highest ADG, followed by FAST, MOD and SLOW. Each category differed from the others in ADG to TW2 (Table 10). While there was a tendency for an effect of strain within category (F_10,106_=1.72; P=0.0849), there were no differences among individual strains within categories for ADG to TW1 or TW2 (adjP>0.34). There was no interaction between target weight and strain within category (F_10,106_=1.08; P=0.38).

### Production and efficiency to TW1 and TW2 by strains within category

There was no difference within CONV or FAST categories for production and efficiency through either of the two target weights (Table 10). For the MOD strains, strain E had higher ADFI through TW1 compared to strain S (Table 6). At TW2, strain O had higher ADFI than strain H. There were no differences among MOD strains in FCR at TW1 or TW2 (Table 10). Through TW1 and TW2, there were no differences among SLOW strains in BW, ADG, ADFI or FCR (Table 10).

### Mortality

Throughout the eight trials, there were no disease outbreaks, and no birds needed treatment with antibiotics or other medications. Overall, mortality rate was 2.52% over the eight trials. There was no relationship between body weight category and overall mortality (Table 11). However, the odds of birds having been culled versus found dead were associated with category. SLOW had greater odds of having fewer culls than all other categories (CONV to SLOW: OR 0.179, CI: 0.041-0.779; FAST to SLOW: OR 0.272, CI: 0.080-0.929; MOD to SLOW: OR 0.134, CI: 0.039-0.466). SLOW had more than twice the odds of having more birds found dead than MOD (OR 2.226, CI: 1.003-4.943).

**Table 11.** Effect of category on percentage of birds culled, found dead and total mortality (raw means ± SE). The odds of having culls and birds found dead differed among categories (P<0.05). The total mortality did not differ among categories. Odds ratios are described in the text.

| Category | Strain | Culls, % | Found dead, % | Total mortality, % |
| --- | --- | --- | --- | --- |
| CONV | | $0.85 \pm 0.27$ | $1.99 \pm 0.55$ | $2.84 \pm 0.63$ |
| FAST | | $0.88 \pm 0.21$ | $1.65 \pm 0.27$ | $2.53 \pm 0.39$ |
| MOD | | $0.80 \pm 0.17$ | $1.47 \pm 0.31$ | $2.27 \pm 0.35$ |
| SLOW | | $0.43 \pm 0.19$ | $2.22 \pm 0.33$ | $2.65 \pm 0.37$ |
| CONV | A<sup>1</sup> | $0.57 \pm 0.57$ | $2.27 \pm 0.93$ | $2.84 \pm 1.09$ |
| | B | $1.33 \pm 0.44$ | $2.46 \pm 0.95$ | $3.79 \pm 1.10$ |
| | C | $0.38 \pm 0.26$ | $1.52 \pm 0.58$ | $1.89 \pm 0.55$ |
| FAST | F | $1.33 \pm 0.52$ | $1.70 \pm 0.49$ | $3.03 \pm 0.64$ |
| | G | $0.76 \pm 0.43$ | $1.14 \pm 0.52$ | $1.89 \pm 0.88$ |
| | I | $0.38 \pm 0.26$ | $1.70 \pm 0.63$ | $2.08 \pm 0.81$ |
| | M | $1.14 \pm 0.43$ | $2.27 \pm 0.43$ | $3.41 \pm 0.74$ |
| MOD | E | $1.33 \pm 0.44$ | $2.27 \pm 0.88$ | $3.60 \pm 0.90$ |
| | H | $0.76 \pm 0.32$ | $0.95 \pm 0.44$ | $1.70 \pm 0.41$ |
| | O | $0.76 \pm 0.32$ | $1.14 \pm 0.59$ | $1.89 \pm 0.78$ |
| | S | $0.38 \pm 0.26$ | $1.52 \pm 0.43$ | $1.89 \pm 0.47$ |
| SLOW | D | $0.19 \pm 0.19$ | $2.08 \pm 0.71$ | $2.27 \pm 0.69$ |
| | J | $0.19 \pm 0.19$ | $2.27 \pm 0.56$ | $2.46 \pm 0.65$ |
| | K | $0.76 \pm 0.43$ | $0.76 \pm 0.32$ | $1.52 \pm 0.43$ |
| | N | $0.57 \pm 0.57$ | $3.79 \pm 0.70$ | $4.36 \pm 0.95$ |
| | T<sup>1</sup> | $2.08 \pm 2.08$ | $1.04 \pm 1.04$ | $3.13 \pm 3.13$ |

<sup>1</sup> Strains A and T were represented in few pens (4) and were therefore not included in statistical analyses for most variables. The data are included for descriptive purposes only.

There were no differences between strains B and C within the CONV category in number of birds culled, number found dead or total mortality. Within the FAST strains, there were significant differences between strains in the number of birds needing to be culled and overall mortality (Table 11). Strain F had more birds needing to be culled than strain I (t_118_=-2.05; P=0.043), had greater overall mortality than strain G (t_117_=−2.14; P=0.0345) and had a tendency for greater mortality than strain I (t_117_=−1.72; P=0.087). Strain M needed to have more birds culled than strain I (t_118_=−2.57; P=0.0115) and tended to have more birds needing to be culled than strain G (t_118_=−1.96; P=0.052). Strain M also had higher total mortality than strain G (t_117_=2.22; P=0.028) and tended to have higher mortality than strain I (t_117_=1.86; P=0.066).

Within the MOD strains, E had higher total mortality than strain O (t_117_=−2.05; P=0.042) and tended to have higher total mortality than strain H (t_117_=−1.70; P=0.091). Within the SLOW strains, strain K had more birds needing to be culled than strain J (t_118_=2.15; P=0.034) and strain D (t_118_=−2.09; P=0.039). Strain N had more birds found dead than strain K (t_117_=3.21; P=0.0017) and tended to have been found dead more than strain D (t_117_=−1.90; P=0.06). Strain J tended to have more birds found dead than strain K (t_117_=−1.96; P=0.053). Strain N tended to have higher total mortality than strain K (t_117_=1.96; P=0.052).

Sudden death syndrome and ascites were rarely observed, and few birds were culled due to lameness. Three of the mortalities were due to sudden death syndrome, whereas 24/54 of the euthanized birds (out of 7,528 birds studied) were euthanized for lameness. Two strains, B (a CONV strain) and F (a FAST strain) accounted for 46% of the culling due to lameness.

## DISCUSSION

There has been increasing interest in the use of slower growing broiler chickens because of their purported improved welfare. Indeed, numerous studies have compared slower growing strains to conventional strains and reported better welfare outcomes for slower growing strains (Nielsen et al., 2003; Bokkers and Koene, 2003, Fanatico et al., 2008; Dixon, 2020; Rayner et al., 2020). Most studies comparing strains have studied them in outdoor, range conditions or compared few strains with disparate growth rates. Our comprehensive study was designed to benchmark welfare and production phenotypes for 16 strains of broiler chickens with a range of growth rates and without outdoor access. Before we could compare the welfare of different strains and determine the relationship between welfare indicators and growth rates, we needed to determine their growth rates and feed efficiencies. The objective of this manuscript was to compare the production, efficiency and mortality of 16 strains of broiler chickens reared in the same environment, under the same management and fed the same diet.

The strains provided to us from commercial breeding companies represented a range of average daily gains from 44 to 68 g/d. We also were provided with one mixed-breed dual-purpose strain with an estimated growth rate of 17.5 g/d. While the strains differed in other ways that may have enabled division, we chose to categorize them based on their growth rates post-hoc to facilitate statistical analyses and generalizations to other strains not studied here. Within categories, strains differed in their growth trajectories and feed efficiencies, which may have had implications on their welfare. We compared categories of strains, and strains within categories at two ‘target weights’-one lighter (estimated at 2.1 kg) and one heavier (estimated at 3.2 kg). We also compared categories and strains at the same age.

The CONV category grew faster, consumed more feed and were more feed efficient than any of the slower growing categories. This was expected as CONV strains of broiler chickens have been selected for fast growth, high feed intake and high feed efficiency over generations (Tallentire et al., 2016). Differences between categories were apparent within the first two weeks of production and became more pronounced as the birds aged. After consumption of similar starter, grower and finisher feed allotments, CONV strains were heavier despite being 6 or more days younger than strains in the other categories. At 48 days of age (when CONV were at TW2 and the others were at TW1), the CONV category was 835-1264 grams heavier than other categories. By the heavier target weight, differences in body weights and feed intake resulted in a 22 to 43-point difference in feed conversion ratios.

Within categories, there were some growth and efficiency differences between strains during early rearing. Differences in growth trajectories to the same final body weight influence physiological and metabolic development (Zubair and Leeson, 1996), which may have impacts on welfare outcomes. Within the CONV category, strains B and C differed in their growth rates through consumption of similar feed allotments. However, differences between the two disappeared by the first target weight. Within the FAST category, strains differed in their early growth and feed intake. Strain M was heavier and had higher early feed intake than other FAST strains, but was lighter and had the worst feed efficiency through consumption of similar feed allotments. As with the CONV category, there were no differences in growth and efficiency within the FAST category by target weight 1. Among the MOD strains, strain E had the highest early ADG and ADFI whereas strain H had the worst early FCR. The higher early ADG for strain E compared to other MOD strains may have affected its mortality rate, as Robinson et al (1992) suggested that early rapid growth contributes to increased metabolic diseases and mortality. Strain H had lower levels of inactivity than other MOD strains through 5 weeks of age (Dawson et al., 2020), which would have influenced feed efficiency. There were differences in ADFI through the heavier target weight, but this did not significantly affect feed conversion ratios. Within the SLOW category through consumption of equal amounts of feed, strain D had the highest feed intake and worst feed conversion ratio; these differences were no longer apparent at the first processing age (48 days).

Mortality within our study was lower than that seen in commercial production systems (National Chicken Council, 2020) and lower than has been reported in some studies comparing fast and slow growing strains (Yalcin et al., 2001; Dixon, 2020; Weimer et al., 2020), but not all (Fanatico et al., 2005). Our flocks were fed antibiotic-free diets but did not experience any disease outbreaks. This may be due to incubation and hatch conditions. We received fertile hatching eggs from our partners and incubated and hatched the eggs at our research facility. This enabled us to vaccinate and place chicks within 8-10 hours, which may have led to improved outcomes compared to other studies (Dixon, 2020; Weimer et al., 2020). There were, however, differences in the rates of culling and being found dead between the slowest growing category and other categories. We observed more early deaths (within the first 10 days) in some of the slowest growing strains, which may indicate yolk sac infections, but the sample size was not large enough to permit statistical analyses. Early mortality may relate to parent-stock variables or hatch conditions (McNaughton et al., 1978; Yassin et al., 2009). All strains were incubated and hatched under the same conditions, to eliminate these factors as confounds. In commercial conditions, incubation and hatch conditions, nutrition and environment are tailored to meet the needs of each genetic strain.

Target weights were chosen to represent typical processing weights for the North American market, and ages for these weights were chosen based on the average days to reach 2.1 and 3.2 kg from breeder estimations. As expected, the CONV strains grew faster, ate more and grew more efficiently than all the remaining strains. Due to project logistics, however, the CONV strains were lighter than the FAST and MOD strains at Target Weight 1. This was due to both the early processing age for CONV (34 d) and the (same) lower density diet used for all strains. We chose to feed all the strains the same three-phase diet, one that had been formulated to meet the needs of a moderate slower growing strain. In one of our pilot studies, we determined that this diet influenced the growth rates of conventional strains (Nascimento dos Santos et al. 2018). However, as we could not formulate individual diets for each strain, we chose to accept this limitation. Similarly, we housed all strains in the same experimental room, with the same room temperature, lighting and management, which may not have been ideal for some, but permitted us to remove these factors as confounds.

When incubated, hatched, housed, managed and fed the same diet, the 16 strains of broiler chickens differed in growth and efficiency, but their growth rates did not influence mortality rates. We categorized strains based on growth rates to a heavy (3.2 kg) weight to enable generalizations based on growth and determine if factors other than growth rates impact the welfare of broiler strains. Determining and categorizing the growth rates of the strains of broiler chickens in this study was the first step in benchmarking their welfare and productivity. Additional manuscripts will discuss the welfare and productivity of these strains, and whether growth rate or other factors influence welfare outcomes for broiler chickens.

## ACKNOWLEDGEMENTS

This work was supported by Global Animal Partnership, Canada First Research Excellence Fund and the in-kind contributions from Ontario Agri-Food Alliance, the anonymous breeding companies and Protekta, Inc. On the processing side, Farm Fresh Poultry, ENS Poultry and Pullets Plus were helpful in ensuring this project ran smoothly. We are grateful for the technical assistance of Cheryl Campbell, Judy Kendall, Sam Leo, Danielle Renwick, Linda Caston and Brian McDougal. This project could not have happened without the diligent help of our student assistants (in alphabetical order by last name): Alan Abdulkadar, Sophia Ahmad, Madeleine Browne, Veronica Cheng, Margaret Daoust, Melanie Felker, King (Stella) Hung, Breanna Jackson, Anna Laszczuk, Narissa Leslie, Siobhan Mellors, Nyasha Mombeshora, Nicole Norena, Quinn Rausch, Erin Ross, Victoria Shouldice, Tara Tanasijevic, Kayley Teal, Priyanka Thavaraja, Megan Weckwerth, Leah Wellard and Jessica Woods. Finally, the assistance of the Arkell Poultry Research Station staff was pivotal to this project’s success. We thank Dave Vandenberg, Innes Wilson, Nancy Wedel, Vern Wideman, Rick Hoiting and Danielle Watson for all their help.

## REFERENCES

AAFC. 2019. Statistics and Market Information, 036. Poultry Production Report by Month/Year; Eviscerated Weight. Accessed Feb. 2019. http://aimis-simia.agr.gc.ca/rp/index-eng.cfm?action=pRr=6pdctc=

Bassler, A.W., C. Arnould, A. Butterworth, L. Colin, I.C. De Jong, V. Ferrante, P. Ferrari, S. Haslam, F. Wemelsfelder, and H.J. Blokhuis. 2013. Potential risk factors associated with contact dermatitis, lameness, negative emotional state, and fear of humans in broiler chicken flocks. Poult. Sci. 92(11):2811–2826.

Beter Leven. 2016. Chicken Standard, 1 Star “VLEESKUIKENS - 1 STER”. Accessed Sept. 2020. https://beterleven.dierenbescherming.nl/app/uploads/sites/2/2020/02/Vleeskuikens-1-ster-Versie-5.1.-ZW-d.d.-01.09.2016.pdf

Bokkers, E.A. and P. Koene. 2003. Behaviour of fast-and slow growing broilers to 12 weeks of age and the physical consequences. Appl. Anim. Behav. Sci. 81(1):59–72.

Bokkers, E.A. and P. Koene. 2004. Motivation and ability to walk for a food reward in fast-and slow-growing broilers to 12 weeks of age. Behav. Proc. 67(2):121–130.

Bokkers, E.A., P.H. Zimmerman, T.B. Rodenburg, and P. Koene. 2007. Walking behaviour of heavy and light broilers in an operant runway test with varying durations of feed deprivation and feed access. Appl. Anim. Behav. Sci. 108(1-2):129–142.

Clune, S., E. Crossin, and K. Verghese. 2017. Systematic review of greenhouse gas emissions for different fresh food categories. J. Clean. Prod. 140:766–783.

Dawkins, M.S. and R. Layton. 2012. Breeding for better welfare: genetic goals for broiler chickens and their parents. Anim. Welf. 21(2):147–155.

Dixon, L.M., 2020. Slow and steady wins the race: The behaviour and welfare of commercial faster growing broiler breeds compared to a commercial slower growing breed. PloS One, 15(4):p.e0231006

Fanatico, A.C., L.C. Cavitt, P.B. Pillai, J.L. Emmert, and C.M. Owens. 2005. Evaluation of slower-growing broiler genotypes grown with and without outdoor access: meat quality. Poult. Sci. 84(11):1785–1790.

Fanatico, A.C., P.B. Pillai, P.Y. Hester, C. Falcone, J.A. Mench, C.M. Owens, C.M. Owens, and J.L. Emmert. 2008. Performance, livability, and carcass yield of slow-and fast-growing chicken genotypes fed low-nutrient or standard diets and raised indoors or with outdoor access. Poult. Sci. 87(6):1012–1021.

Federal Office for Agriculture and Food, Working Group of the Federal States on Organic Farming 2009. Auslegung der Rechtsvorschriften für den Ökolandbau. Accessed Sept. 2020. https://www.oekolandbau.de/service/rechtsgrundlagen/auslegungen-der-eu-rechtsvorschriften/protokoll-details/?tx_oekolbloek_loekdetail%5BsessionUid%5D=56tx_oekolbloek_loekdetail%5BagendaItemUid%5D=340tx_oekolbloek_loekdetail%5Baction%5D=detailtx_oekolbloek_loekdetail%5Bcontroller%5D=LoekcHash=48be4c5187e463ae78488b85c0847d66.

Hartcher, K.M. and H.K. Lum. 2020. Genetic selection of broilers and welfare consequences: a review. World’s Poult. Sci. J. 76(1):154–167.

Havenstein, G.B., P.R. Ferket, and M.A. Qureshi. 2003. Growth, livability, and feed conversion of 1957 versus 2001 broilers when fed representative 1957 and 2001 broiler diets. Poult. Sci. 82(10):1500–1508.

Kapell, D.N.R.G., W.G. Hill, A.M. Neeteson, J. McAdam, A.N.M. Koerhuis, and S. Avendano. 2012a. Twenty-five years of selection for improved leg health in purebred broiler lines and underlying genetic parameters. Poult. Sci. 91(12):3032–3043.

Kapell, D.N.R.G., W.G. Hill, A.M. Neeteson, J. McAdam, A.N.M. Koerhuis, and S. Avendano. 2012b. Genetic parameters of foot-pad dermatitis and body weight in purebred broiler lines in 2 contrasting environments. Poult. Sci. 91(3):565–574.

Kittelsen, K.E., B. David, R.O. Moe, H.D. Poulsen, J.F. Young and E.G. Granquist. 2017. Associations among gait score, production data, abattoir registrations, and postmortem tibia measurements in broiler chickens. Poult. Sci. 96(5):1033–1040.

McGeown, D., T.C. Danbury, A.E. Waterman-Pearson, and S.C. Kestin. 1999. Effect of carprofen on lameness in broiler chickens. Vet. Rec. 144(24):668–671.

McNaughton, J.L., J.W. Deaton, F.N. Reece, and R.L. Haynes. 1978. Effect of age of parents and hatching egg weight on broiler chick mortality. Poult. Sci. 57(1):38–44.

Mohammadigheisar, M., R.B. Shirley, J. Barton, A. Welsher, P. Thiery, and E. Kiarie. 2019. Growth performance and gastrointestinal responses in heavy Tom turkeys fed antibiotic free corn− soybean meal diets supplemented with multiple doses of a single strain Bacillus subtilis probiotic (DSM29784). Poult. Sci. 98(11):5541–5550.

Mohammadigheisar, M., V.L. Shouldice, S. Torrey, T. Widowski, and E.G. Kiarie. 2020. Research Note: Comparative gastrointestinal, tibia, and plasma attributes in 48-day-old fast-and slow-growing broiler chicken strains. Poult. Sci. 99(6):3086–3091.

Nääs, I.A., I.A. Paz, M.S. Baracho, A.G. Menezes, L.G.F. Bueno, I.C.L. Almeida, and D.J. Moura. 2009. Impact of lameness on broiler well-being. J. Appl. Poult. Res. 18(3):432–439.

Nascimento dos Santos, M., T. Widowski, E. Kiarie, M. Mohammadigheisar, I. Mandell and S. Torrey. 2018. The impact of low-nutrient and conventional diets on lameness, tibial dimensions and tibial ash concentrations for two fast-growing genotypes of broiler chickens. Poult. Sci. 97(e-suppl 1):166.

National Chicken Council. 2020. US Broiler Performance, 1925 to present. Accessed Sept. 2020. https://www.nationalchickencouncil.org/about-the-industry/statistics/u-s-broiler-performance/

National Farm Animal Care Council, 2016. The Code of Practice for the Care and Handling of Chickens, Turkeys and Breeders. Accessed Jan. 2020. https://www.nfacc.ca/pdfs/codes/poultry_code_EN.pdf

Nielsen, B.L., M.G. Thomsen, P. Sørensen, and J.F. Young. 2003. Feed and strain effects on the use of outdoor areas by broilers. Br. Poult. Sci. 44(2):161–169.

Opengart, K., S.F. Bilgili, G.L. Warren, K.T. Baker, J.D. Moore, and S. Dougherty. 2018. Incidence, severity, and relationship of broiler footpad lesions and gait scores of market-age broilers raised under commercial conditions in the southeastern United States. J. Appl. Poult. Res. 27(3):424–432.

Rayner, A.C., R.C. Newberry, J. Vas, and S. Mullan. 2020. Slow-growing broilers are healthier and express more behavioural indicators of positive welfare. Sci. Rep. 10(1):1–14.

Rekaya, R., R.L. Sapp, T. Wing, and S.E. Aggrey. 2013. Genetic evaluation for growth, body composition, feed efficiency, and leg soundness. Poult. Sci. 92(4):923–929.

Robinson, F.E., H.L. Classen, J.A. Hanson, and D.K. Onderka. 1992. Growth performance, feed efficiency and the incidence of skeletal and metabolic disease in full-fed and feed restricted broiler and roaster chickens. J. Appl. Poult. Res. 1(1):33–41.

Tallentire, C.W., I. Leinonen, and I. Kyriazakis. 2016. Breeding for efficiency in the broiler chicken: A review. Agron. Sustain. Dev. 36(4):66.

Volaille Label Rouge, 2020. A broad range of Label Rouge Poultry. Chicken. Accessed Sept. 2020. http://www.volaillelabelrouge.com/en/chicken/

Weimer, S.L., A. Mauromoustakos, D.M. Karcher, and M.A. Erasmus. 2020. Differences in performance, body conformation, and welfare of conventional and slow-growing broiler chickens raised at 2 stocking densities. Poult. Sci. 99(9):4398–4407.

Wilhelmsson, S., J. Yngvesson, L. Jönsson, S. Gunnarsson, and A. Wallenbeck. 2019. Welfare Quality^®^ assessment of a fast-growing and a slower-growing broiler hybrid, reared until 10 weeks and fed a low-protein, high-protein or mussel-meal diet. Livestock Sci. 219:71–79.

Yalcin, S., S. Özkan, L. Türkmut, and P.B. Siegel. 2001. Responses to heat stress in commercial and local broiler stocks. 1. Performance traits. Br. Poult. Sci. 42(2):149–152.

Yassin, H., A.G. Velthuis, M. Boerjan, and J. van Riel. 2009. Field study on broilers’ first-week mortality. Poult. Sci. 88(4):798–804.

Zubair, A.K. and S. Leeson. 1996. Compensatory growth in the broiler chicken: a review. World’s Poult. Sci. J. 52(2):189–201.

Zuidhof, M.J., B.L. Schneider, V.L. Carney, D.R. Korver, and F.E. Robinson. 2014. Growth, efficiency, and yield of commercial broilers from 1957, 1978, and 2005. Poult. Sci. 93(12):2970–2982.

